# Increased grain weight conferred by *GW2* mutations in wheat does not translate into yield gains in multi-year field trials of near-isogenic lines

**DOI:** 10.64898/2025.12.04.692284

**Authors:** James Simmonds, Pamela Crane, Sophie Eade, Aura Montemayor-Lara, Matt Kerton, Nicholas Bird, Phil Tailby, Peter Jackson, Duncan Warner, Charlotte Hayes, David Schafer, Cristobal Uauy

## Abstract

Multiple studies have identified genes affecting grain morphology, yet their capacity to deliver yield gains under field conditions remains unclear. We performed a multiyear, multilocation factorial evaluation of *GRAIN WIDTH2* (*TaGW2*) mutants in hexaploid wheat using BC_4_ near-isogenic lines, sowing-density treatments and semi-dwarfing *RHT1* backgrounds. Loss-of-function mutations in *TaGW2* increased grain size and thousand grain weight (TGW) additively; with the *aaBBDD* single mutant showing the most stable single-locus effect, while the *aabbdd* triple mutant achieved ~20% higher TGW across twelve field trials. However, overall grain yield remained unchanged or slightly reduced, reflecting a compensatory trade-off with grain number. Spike phenotyping of both main and secondary tillers showed comparable increases in TGW and spike yield despite fewer grains per spike, indicating that limited yield gain primarily reflects reduced spike number per unit area rather than decreased spike-level productivity. Effects were stable across sowing densities, whereas interactions with semi-dwarfing alleles were allele-specific: *RHT-B1b* partially suppressed TGW gains and accentuated yield penalties, whereas *RHT-D1b* maintained the large-grain phenotype and productivity. Across experiments, the *TaGW2-A1D1* double mutant increased TGW (~14%) while maintaining yield stability, identifying it as a promising genotype for breeding. We conclude that *TaGW2* is a reliable modifier of grain size but not yield in isolation.

## Introduction

Wheat (*Triticum aestivum* L.) is a staple crop providing a major source of calories and protein for the global population. As global wheat demand continues to outstrip the rate of yield improvement (FAO, 2018; Fischer et al., 2014), future food security faces considerable challenges, especially under ever increasing environmental pressures. Yield improvements are also at the heart of initiatives to reduce greenhouse gas emissions in agriculture; for example, the UK Seventh Carbon Budget calls for a 16% increase in crop yields as a key driver to half CO_2_ emissions by 2050 (Climate Change Committee, 2025). Accordingly, the identification, functional characterization, and incorporation of genes and alleles that enhance yield represent key priorities for breeding programs worldwide.

Grain yield in wheat is a complex polygenic trait, determined primarily by its three major components: the number of spikes per area, the number of grains per spike and the average weight of individual grains. Wheat yield improvements have traditionally relied on optimising grain number per unit area, achieved through the combined effects of genetic selection and enhanced agronomic practices (Calderini et al., 1995; Slafer et al., 2021). More recently there have been numerous studies that propose grain size and weight as a method of yield improvement (Adamski et al., 2021; Brinton et al., 2017; Calderini et al., 2021; Simmonds et al., 2014). The rationale for focussing efforts on gene discovery for grain morphometrics is that grain size is the final yield component to be determined, is stably inherited and has higher heritability than crop yields.

One gene that has attracted significant attention is *GRAIN WIDTH 2* (*GW2*), a RING-type E3 ubiquitin ligase that negatively regulates grain size in rice (Song et al., 2007) and whose function seems conserved across cereals (Li et al., 2010; Long et al., 2024; Tao et al., 2017). Wheat orthologs of *GW2* (*TaGW2*) have been shown to play similar roles, with targeted loss of function mutations resulting in larger grains and increased thousand grain weight (TGW) by promoting cell expansion in the grain pericarp (Simmonds et al., 2016; Zhang et al., 2018). Previously we demonstrated in BC_1_F_3_ lines grown in controlled environment rooms that mutations from the three wheat homoeologues act in an additive manner, with grain size effects ranging from ~5.5% in the single mutants, 13.4% in the double mutants and 20.7% in the full knock-out (triple) (Wang et al., 2018). Subsequently, additional studies confirmed these effects in CRISPR induced mutants (Achary & Reddy, 2021; Kis et al., 2024; Zhang et al., 2018).

However, the effect of *TaGW2* (referred to hereafter as *GW2*) mutations on final yield remains context-dependent and may vary across genetic backgrounds and environments due to compensatory effects on grain number and source-sink dynamics (Vicentin & Calderini, 2025). The extent to which these improvements translate into consistent yield advances are rarely reported, especially from field-based yield experiments, across multiple sites and years. This study aims to evaluate the relationship between grain weight and yield in *GW2* near-isogenic lines (NILs) across multiple field environments and years. We utilised BC₄ NILs developed from wheat *GW2* TILLING mutations in a UK elite genetic background (Simmonds et al., 2016; Wang et al., 2018) to assess the gene’s phenotypic effects while minimizing confounding variation from unrelated genomic regions. We compared the effects of individual *GW2* homoeologues as well as their combined contributions, providing insights into the additive and potentially compensatory roles of these genes in wheat yield determination. In addition, we evaluated the effects of *GW2* homoeologs under contrasting *REDUCED HEIGHT 1* (*RHT1*) alleles, given that semi-dwarfing *RHT1* alleles are known to negatively affect grain size but have positive effects on yield and grain number (Flintham et al., 1997; Jobson et al., 2018; Keyes & Sorrells, 1989). Through understanding the pleiotropic effects of *GW2* alleles on yield components, we can assess the stability of *GW2*associated phenotypes and their implications for wheat yield improvement strategies.

## Materials & Methods

### Plant Materials

We previously identified mutations in the three *GW2* homoeologues from the tetraploid *Triticum turgidum* cv Kronos and hexaploid *T. aestivum* cv Cadenza TILLING populations (Krasileva et al., 2017; Simmonds et al., 2016; Wang et al., 2018) (Supplemental Table 1). Mutations were individually crossed to cv Paragon and the F_1_ then intercrossed to combine all mutations into a single plant. The presence of mutations was confirmed through marker assisted selection with SNP specific KASP markers (Supplementary Table 2). Heterozygous triple mutant plants (*AaBbDd*) were self-pollinated to produce homozygous BC_1_F_2_ plants as described in Wang et al., 2018. The homozygous BC_1_ plants were backcrossed again and BC_2_F_1_ plants were self-pollinated and homozygous wildtype and mutant plants for all homoeologue combinations were extracted at BC_2_F_2_. The BC_2_ NILs were utilised in field Experiment 1 (2018). Backcrossing continued and advanced to BC_4_F_2_ NILs to minimise offtarget regions and these were sown in Experiments 2-5.

As Paragon does not carry either of the main *RHT1* semi-dwarfing alleles utilised in the UK (*RHT-D1b*, *RHT-B1b*) we wanted to assess the interaction of the full *GW2* knock-out with these alleles. BC_4_ *GW2* triple mutants NILs were crossed to Paragon *RHT-D1b and* Paragon *RHT-B1b* NILs, developed as part of the ‘Paragon Library’ through the Wheat Genetic Improvement Network (WGIN). The F_1_ seed were self-pollinated and after screening with KASP markers, homozygous F_2_ lines with the *GW2* triple mutant alleles or three wildtype alleles, and with either *RHT-D1b, RHT-B1b or* wildtype alleles at *RHT1*, were selected. For all genotypes we developed two independent crossing streams. Analyses, however, were performed by genotype and therefore streams were combined.

Seed of the Paragon BC_4_ *GW2* single homoeologous mutants and the Paragon BC_4_ *GW2* triple mutant used in this study are available via the JIC Germplasm Resources Unit (https://seedstor.ac.uk/) with the following accession numbers WM0011 (A genome), WM0005 (B genome), WM0007 (D genome), WM0015 (triple mutant). Seed of the Paragon *RHT* NILs and Paragon *GW2*RHT* NILs are also available to order; WParNil0077 (Paragon *RHT-D1b*), WParNil0078 (Paragon *RHT-B1b)*, WM0325 (Paragon *GW2_triple + RHT-D1b*) and WM0326 (*GW2_triple + RHT-B1b*).

### Field Experiments

All field trials, apart from Experiment 4, were sown at The Morley Agricultural Foundation, Wymondham, Norfolk, UK (52.55421, 1.03274). BC_2_ / BC_4_ NILs were sown in the Autumn in 2018 and 2020 (Exp. 1 & 3) respectively, but in early Spring in 2020 (Exp.2) due to unfavourable weather/soil conditions in the Autumn 2019. Experiments 1, 2, 3, 5 and 6 were sown in yield plots 6 m x 1.2 m aiming for a target population of 275 plants*m^−2^, consisting of 8 rows with 0.13 m spacing. Trials were sown in a randomised complete block design with either 3 blocks (Exp.1) or 5 blocks (Exp. 2, 3, 5 and 6) and with two or three NILs for each genotype developed from independent crossing streams. Plot ends were removed pre-harvest to leave a 4 m x 1.2 m combinable plot area.

In 2020 the BC_4_ NILs of the triple mutant (*aabbdd*) and wildtype (*AABBDD*) were also sown in multi-site trials as part of the ‘Designing Future Wheat’ Breeders Toolkit (Exp 4). These were sown in three replicate trials at multiple locations across England with the breeding companies DSV, Elsoms, KWS, Limagrain, NPZ UK and Syngenta.

To test performance of the *GW2* BC_4_ NILs at different planting densities, we sowed a trial in 2022 (Exp.5) at three seed rates, the standard 275 plants*m^−2^, an extremely high seed rate of 500 plants*m^−2^ and a low seed rate aiming for 125 plants*m^−2^.

To test if *GW2* interacted with *RHT1* we developed *GW2*RHT-B1b/D1b* NILs and these were also sown in 2022 at the standard seed density (275 plants*m^−2^). All trials listed above received standard farm pesticide and fertiliser applications to reproduce commercial practise. The locations and details of all field experiments are highlighted in Supplementary Table 3.

### Phenotypic Assessments

The following phenotypic assessments were made, with corresponding Crop Ontology classifications and a full breakdown of traits scored in each experiment provided in Supplementary Table 4. Heading date was assessed as the number of days from 1^st^ May until ¾ of the canopy had reached ¾ emergence (Zadocks growth stage 57) (Zadoks et al., 1974). Maturity was scored when the green colour in the peduncles had been lost in 50% of the plot and calculated as the number of days from 1^st^ May until maturity. Grain Filling Period was calculated as the difference in days between Heading and Maturity. Tiller number was assessed by counting the number of fertile tillers along two 1 m rows per plot and calculating the average, always using the second row from each side of the plot. Crop height was measured from the soil level to the average height of the canopy.

For Experiments 1-3, prior to harvest, ten main spikes were sampled from each plot for spike yield components including spikelet number, rudimentary basal spikelets, seeds/spike, yield/spike and seed/spikelet. For Experiments 5 and 6, two 1 m strips were marked out prior to tiller counts to ensure the same area was counted at both growth stages. These sections were then hand-pulled from each plot to enable the analysis of the above spike yield components to be carried out separately for main tiller and secondary tillers. After separation, all main spikes and secondary spikes were counted and a ratio of secondary spikes to main spikes was calculated. Of these, we randomly sampled five main spikes and five secondary spikes from each 1 m strip to analyse for spike yield components (ten total spikes per plot). All plots were combined and the following measurements recorded: plot yield, grain moisture and hectolitre weight. Post-harvest grain samples were taken from each plot and morphometric measurements (grain width, length, area and thousand grain weight) were recorded from ~500 grains per sample using the MARViN grain analyser (GTA Sensorik GmbH, Germany). In addition, seed was analysed for protein, starch, fibre, dry gluten content and grain hardness using the Perten DA 7250 NIR Analyser (PerkinElmer, Perten Instruments, Hägersten, Sweden).

All raw data from Experiments 1-6 have been deposited on Zenodo (Simmonds., 2025).

### Statistical Analysis

Experiments 1–3 (2019–2021) were analysed separately using fixed-effects ANOVA models of the form *Trait ~ Genotype + Block*, with both factors treated as fixed. Tukey’s Honest Significant Difference (HSD) test (α = 0.05) was used for post hoc comparisons among genotypes. To assess overall effects across years, a combined two-way ANOVA (*Trait ~ Genotype + Year + Genotype × Year*) was performed to evaluate main effects and interactions.

For Experiment 4, data from multiple trial locations were analysed both collectively (*Trait ~ Genotype + Location + Genotype × Location*) and separately within each site using one-way ANOVA models (*Trait ~ Genotype + Block*).

In Experiment 5, effects of genotype, seed rate, and their interaction were tested using a multi-factor ANOVA (*Trait ~ Genotype + SeedRate + Block + Stream + Genotype × SeedRate*). Tukey’s HSD test identified significant differences among genotypes and seed rates. An additional one-way ANOVA compared all genotype × seed rate combinations (factor: *GW2_SR*).

For Experiment 6, a two-way ANOVA (*Trait ~ Rht_Allele + GW2_ABD + Block + Rht_Allele × GW2_ABD*) assessed main and interaction effects, followed by Tukey’s HSD test. A separate one-way ANOVA examined the combined effects of *RHT_GW2* with replication included.

All analyses were conducted in R using base functions (aov) (RStudio Team, 2025) and the HSD.test() function from the agricolae package (Mendiburu, 2019).

## Results

### Individual GW2 homoeologues confer modest increases in grain size and weight

All combinations of the BC_4_ single, double and triple mutant NILs were evaluated in three field trials over consecutive years (2019-2021; Experiments 1-3) to assess their effects on grain size, grain weight, final grain yield, and yield components.

Among the single mutants, *GW2_A1* (*aaBBDD*) was the highest performing, increasing thousand grain weight (TGW) by an average of 6.7% relative to the wildtype (WT) control NIL (range 4.4-7.4%; Figure 1; Supplementary Table 5), consistent with previously reported results (Simmonds et al., 2016). Although only significant in 2020, combined analysis across all three trials indicated that *GW2_A1* was the only single mutant with a significant benefit on TGW. This was due to both significant increases in grain width and length. *GW2_B1* (*AAbbDD*) also exhibited significantly increased grain length in 2019 and in the overall analysis, but its 1.1% increase in yield compared to WT was non-significant. *GW2_D1* (*AABBdd*) showed no significant differences in TGW or grain size parameter compared to WT, supporting previous findings (Wang et al., 2018).

**Figure 1.**
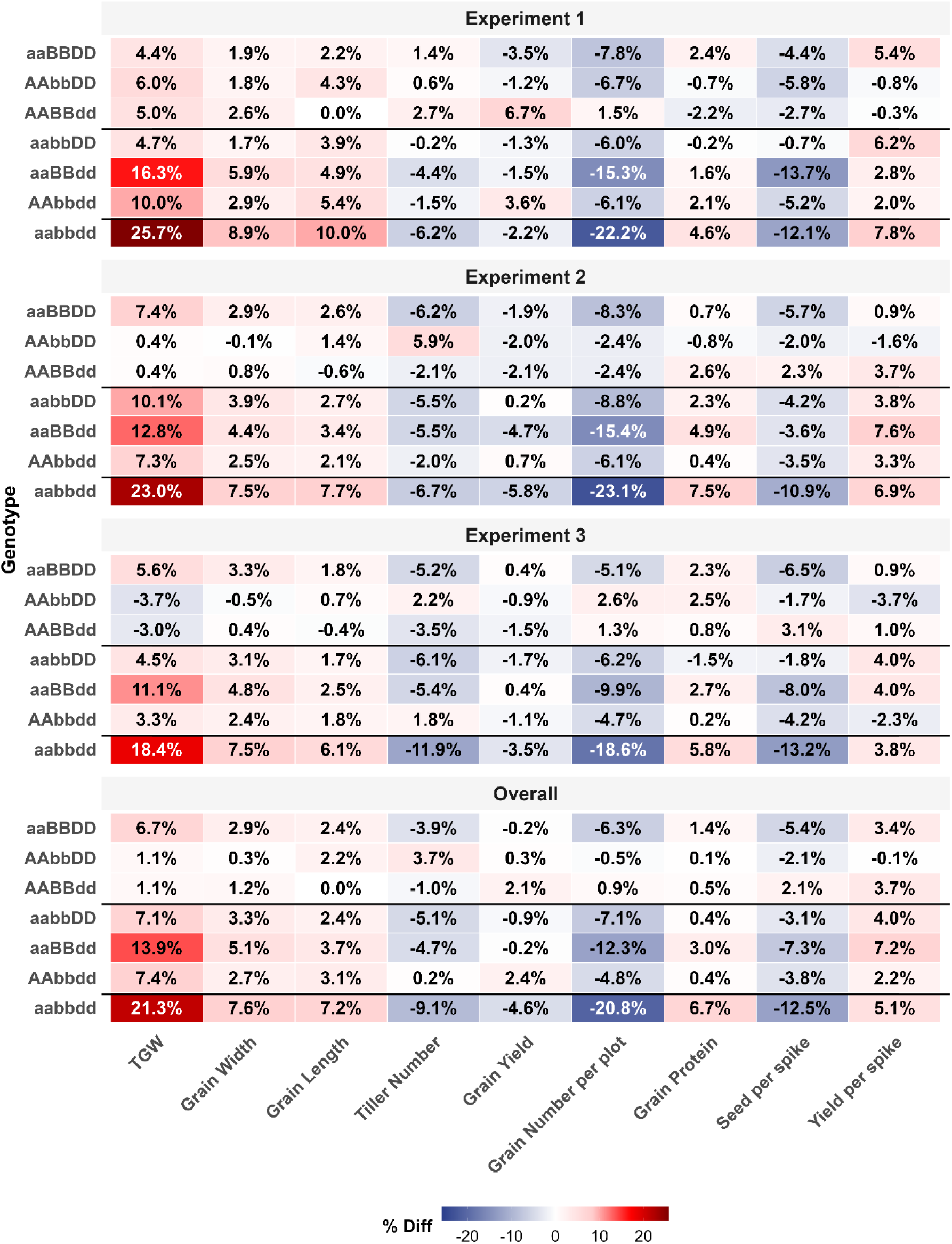
Heatmap showing the percentage difference of single (aaBBDD, AAbbDD, AABBdd), double (aabbDD, aaBBdd, AAbbdd), and triple (aabbdd) *GW2* mutant NILs relative to the wild-type control NIL in Experiments 1-3 (2019-2021) and the overall analysis. Positive differences are shown in increasing intensity of red, while negative differences are shown in increasing intensity of blue. Statistical significance is not displayed on the heatmap but is provided in Supplementary Table 4.

For other traits, the only significant differences from WT were observed for *GW2_A1*, which showed reduce calculated Grain Number per plot (cGNPlot) in 2020 and reduced cGNPlot and seeds per spike in the overall analysis. None of the single mutants showed a significant effect on overall grain yield (Figure 1; Supplementary Table 5).

### GW2 homoeologs act additively to increase grain size and weight (Experiments 1-3)

Combinations of *GW2* mutants were assessed for their combined effect on productivity traits across the three field trials (Experiment 1-3). For TGW, all double mutants (*aabbDD*, *aaBBdd* and *AAbbdd*) showed significant increases compared to the WT. Among them, *GW2_A1D1* (*aaBBdd*) had the largest and most consistent effect, with TGW increases in all three experiments and an overall increase of 13.9% compared to ~7% for the other double mutant combinations. The triple mutant *GW2_A1B1D1* (*aabbdd*) showed a further increase, with TGW 21.3% higher than WT (*P* < 0.001) and significantly higher than any doublemutant (Figure 1; Supplementary Table 5). This increase was driven by proportional increases in both grain width and length, consistent with previous findings (Wang et al., 2018). Despite these marked increases in grain size and weight, grain yield in the double mutants did not differ significantly from the WT. Analysis of yield components revealed compensatory reductions on a series of traits that contribute to yield, including final tiller number (−0.2% to –5.1%), grains per plot (–4.8% to –12.3%), and seeds per spike (–3.1% to –7.3%). These negative effects were further exacerbated in the triple mutant, resulting in a significant 4.6% reduction in grain yield relative to the WT NIL (Figure 1; Supplementary Table 5). Across all measured traits, the triple mutant showed significant increases in TGW (21.3%), grain width (7.6%), grain length (7.2%), grain protein concentration (6.7%) and dry gluten content (7.9%). However, several yield component traits were significantly decreased, including tiller number (−9.1%), hectolitre weight (−2.0%), grains per plot (−20.8%), spikelet number (−3.4%), seeds per spike (−12.5%), and seeds per spikelet (−8.6%) (Figure 1; Supplementary Table 5).

Interestingly, despite the triple mutant producing spikes with fewer seeds, the total grain weight per spike (yield per spike) was consistently higher than WT, ranging from 3.8 to 7.8%, although the differences were not significant. These findings highlight a trade-off between grain size and yield components, which motivated further experiments (Experiments 5 and 6) to dissect the basis of this relationship.

### Multi-Site Breeders trials confirm consistent effects of GW2 mutants (Experiment 4)

To validate the results from the multi-year, single site trials, we conducted a multi-site trial at six UK wheat breeders field stations across England (Experiment 4) comparing the *GW2_A1B1D1* triple mutant (*aabbdd*) and the wildtype (*AABBDD*). Data were analysed both for individual sites and across all locations (Supplementary Table 6). TGW was significantly increased at all sites, with an average gain of 18.2% relative to WT, confirming the robust and stable effect of *GW2* on grain weight observed in Experiments 1-3. Grain protein concentration (4.6%) and dry gluten content (8.8%) were also significantly higher in the triple mutant (supporting the data from Experiments 1-3) and showed consistent improvement across locations, although not all sites were statistically significant. Across all sites, grain yield was unchanged between genotypes (8.912 T*ha^−1^ for the triple mutant versus 8.905 T*ha^−1^ for WT; *P* = 0.971). Yield was significantly decreased at one of site (Elsoms −9.6%; *P* = 0.02), whereas in all other locations the triple mutant gave slight, nonsignificant yield increases (ranging from 0.5 to 7.8%). Based on the yield and TGW values, the calculated grain number was significantly reduced by 15.4% across environments. Thus, the multi-site breeder trials confirmed that *GW2*-mediated increases in grain size and weight are stable across diverse UK environments, but yield remains unchanged due to a consistent trade-off with grain number.

### Grain size and yield are unaffected by sowing density (Experiment 5)

To further investigate the nature of this trade-off, we grew the *GW2-A1B1D1* triple mutant and wildtype NILs at three sowing densities: low (125 plants*m^−2^), standard rate (275 plants*m^−2^) and high (500 plants*m^−2^) in Experiment 5. This experiment was designed to manipulate tillering potential, promoting high tillering in the low seed rate and reduced tillering in the high seed rate, and to evaluate whether *GW2* effects on productivity were influenced by plant density. Previously we observed that tillers*m^−1^ assessed pre-harvest (GS87) were consistently lower in the triple mutant in all three experiments, and 9.1% lower on average (Figure 1; Supplementary Table 5). For this experiment, we additionally measured plant establishment, tiller number at GS32 (just prior to tiller abortion) and calculated the tillers per plant at each stage and the rate of tiller abortion.

Across genotypes, sowing density had the expected, significant effect on plant establishment: low, standard and high seed rate produced 22.2, 32.4, and 45.6 plants*m^−1^, respectively (Supplementary Table 7). Tiller production was also successfully manipulated, with tiller numbers per plant of 3.3 (low), 1.9 (standard) and 1.4 (high).

Interestingly, we discovered that the triple mutant exhibited reduced plant establishment at all three seed rates, significantly so at the high density (Figure 2A; Supplementary Table 7). At GS32, tiller counts were significantly lower in the triple mutant at the high seed rate (119.2 tillers*m^−1^ versus 133.0 tillers*m^−1^ for WT), but not at the standard sowing rate. At preharvest, the expected reduction in tiller numbers was evident at both the standard (−13%) and high (−10%) seed rates, but not at the low seed rate. Despite evidence of some subtle interactions (Figure 2A), tiller abortion rates were unaffected by *GW2* (~43% in both genotypes), and increased significantly with higher sowing densities.

**Figure 2.**
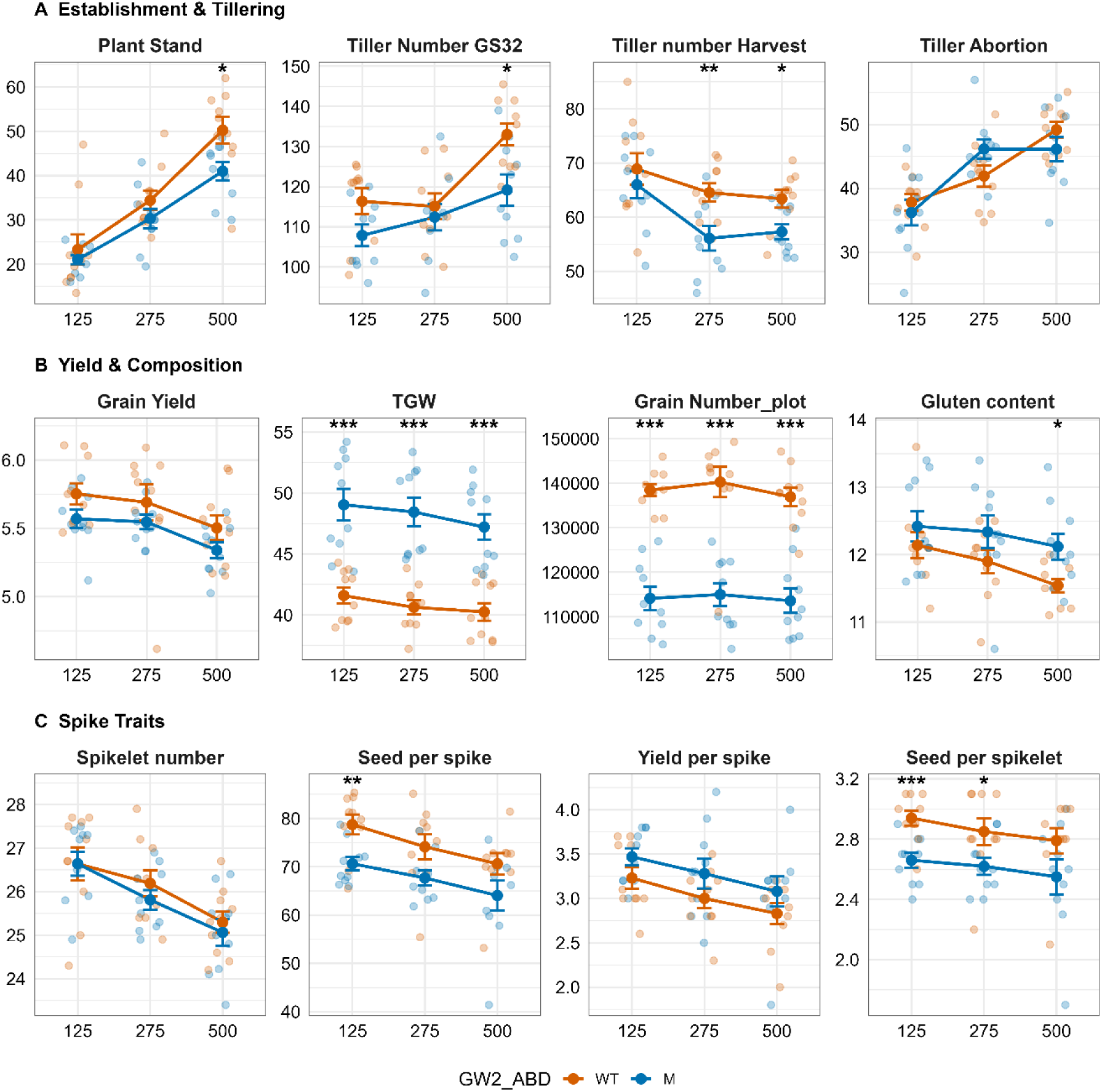
Interaction plots of selected traits for BC_4_ *GW2* NILs sown at three seed densities (125, 275, 500 plants*m^−2^) in Experiment 5. Wildtype NILs are shown in orange and the *GW2* triple mutant NILs in blue. Mean values (solid circles) are presented with standard error bars. Significance levels between WT and triple mutant (M) NILs at each seed rate were determined by two-sample t-tests: *** *P* < 0.001, ** *P* < 0.01, * *P* < 0.05; no symbol indicates *P* ≥ 0.05. Panels represent groups of traits: (A) establishment and tillering (recorded from two 1 m rows per plot), (B) yield and grain composition, and (C) spike traits (measured from ten sampled main spikes).

Seed rate had little impact on the effect of *GW2* on yield and grain quality traits (Figure 2B). TGW was significantly increased in the triple mutant at all three seed rates (mean 18.2%), while grain yields were significantly lower when analysed across all seed rates (−2.9% relative to WT). Interestingly, within genotypes, yields were similar at the low and normal seed rates, but significantly lower at the high density. Spike yield components were also unaffected by seed rate (Figure 2C); consistent with previous experiments, the triple mutant produced fewer seeds per spike and seeds per spikelet, but overall increased yield per spike from main spikes. Taken together, these results indicate that *GW2* effects on grain size and yield are stable across sowing densities, with no evidence of genetic interaction between *GW2* allelic status and plant density.

### No compensatory effect from secondary tillers (Experiment 5)

Analysis of spike yield components indicated that in main spikes, mutations in *GW2* led to an inverse relationship between grain size and grain number yet yield per spike was consistently higher. However, this advantage was not reflected in total plot yield. We therefore asked whether our focus on main spikes might have biased the results, and whether secondary tillers might compensate for reduced tiller number or altered spike traits. Given that tillering was experimentally manipulated by sowing density, we expected a higher proportion of secondary spikes to main spikes at low density and fewer at higher density.

To test this, we phenotyped ten main spikes and ten secondary spikes per plot for both genotypes across the three sowing densities. Analysis of the percentage difference between the triple mutant and WT revealed no significant differences in the response of main versus secondary spikes for any major spike yield component (Figure 3A; Supplementary Table 7). The relative percentage difference between *GW2* alleles were highly consistent across spike types and seed rates: both main and secondary spikes exhibited higher TGW, fewer seeds per spike and per spikelet, and overall higher spike yield compared to WT.

**Figure 3.**
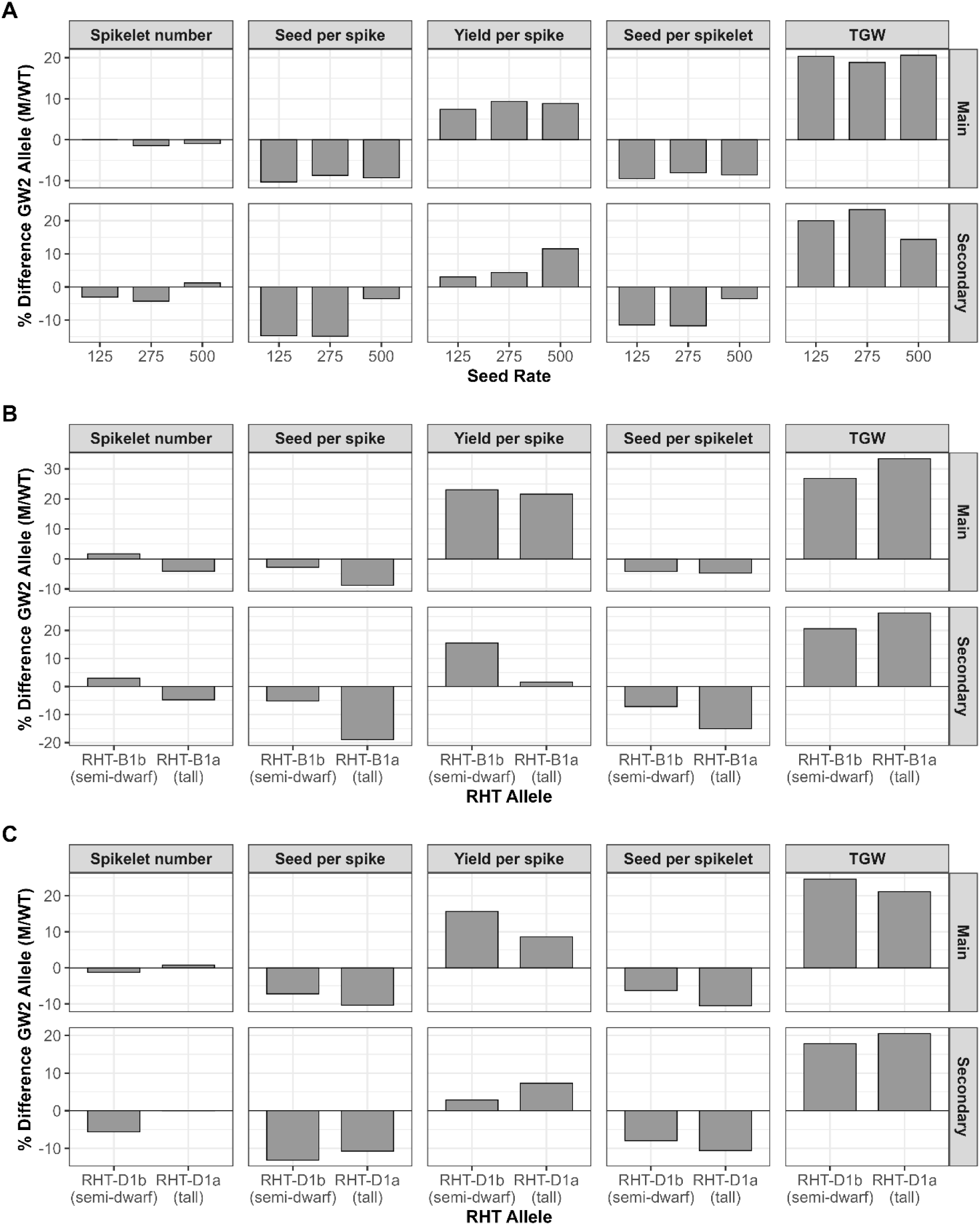
Divergent bar charts showing the percentage difference between *GW2* mutant and WT NILs for selected traits. (A) BC_4_ *GW2* NILs from Experiment 5; (B) BC_3_ *GW2*RHT-B1* NILs from Experiment 6, and (C) BC_3_ *GW2*RHT-D1* NILs from Experiment 6. Bars represent the percentage difference between triple mutant and WT NILs for both main and secondary spikes across seed rates (A) or *RHT* allele background (B, C). Two-way ANOVA was performed for each trait to test for interactions between spike type (main vs secondary) and genotype. No significant differences were detected (*P* ≥ 0.05) for any trait.

As intended, the ratio of secondary to main spikes (ST_MT_EarN_r) varied significantly with seed rate; 2.3, 1.0 and 0.5 secondary spikes per main spike at the low, standard and high densities, respectively, but showed no interaction with *GW2* (*P* = 0.758). This indicates that the main and secondary spikes responded similarly. Although the magnitude of effects was slightly smaller in secondary spikes, the overall yield per spike still increased, suggesting that the reduction in total grain yield in *GW2* mutants primarily results from fewer tillers or spikes, rather than from altered spike-level productivity.

### GW2 phenotype is unaffected by RHT1 (Experiment 6)

Given that the parental background (cv. *Paragon*) carries the wild-type alleles at both major dwarfing loci (*RHT-B1a* and *RHT-D1a*), we developed BC₃ *GW2 × RHT-B1/D1* near-isogenic lines and evaluated them in Experiment 6. Previous studies have demonstrated that *RHT* alleles typically reduce grain size but increase grain number, thereby enhancing overall yield (Flintham et al., 1997; Jobson et al., 2018; Keyes & Sorrells, 1989). Conversely, *GW2* mutations decrease grain number and increase seed size. This experiment therefore provided an opportunity to investigate the potential interaction between these contrasting genetic effects.

As expected, crop height was significantly reduced in lines carrying *RHT-B1b* (−11%) and *RHT-D1b* (−15%), confirming the presence of the semi-dwarfing alleles (Supplementary Tables 8_A and 8_B).

For thousand grain weight, we observed the expected reductions in both *RHT-B1b* (Figure 4_A; Supplementary Table 8_A) and *RHT-D1b* (Figure 4_B; Supplementary Table 8_B) backgrounds compared to WT, consistent with the known effects of *RHT* on grain size, with *GW2* performing significantly worse in the presence of *RHT_B1*. When analysed independently of *GW2* status, TGW decreased significantly by 11% and 6% for *RHT-B1b* and *RHT-D1b*, respectively. However, a significant interaction was observed between *RHT-B1* and *GW2* (*P* < 0.02): the *GW2* triple mutant *× RHT-B1a* displayed a significant TGW increase when in a *RHT-B1b* semi-dwarf background. No such interaction was detected with *RHT-D1b*, where TGW differences between *GW2* mutant and WT lines were proportionally similar regardless of *RHT-D1* status.

**Figure 4.**
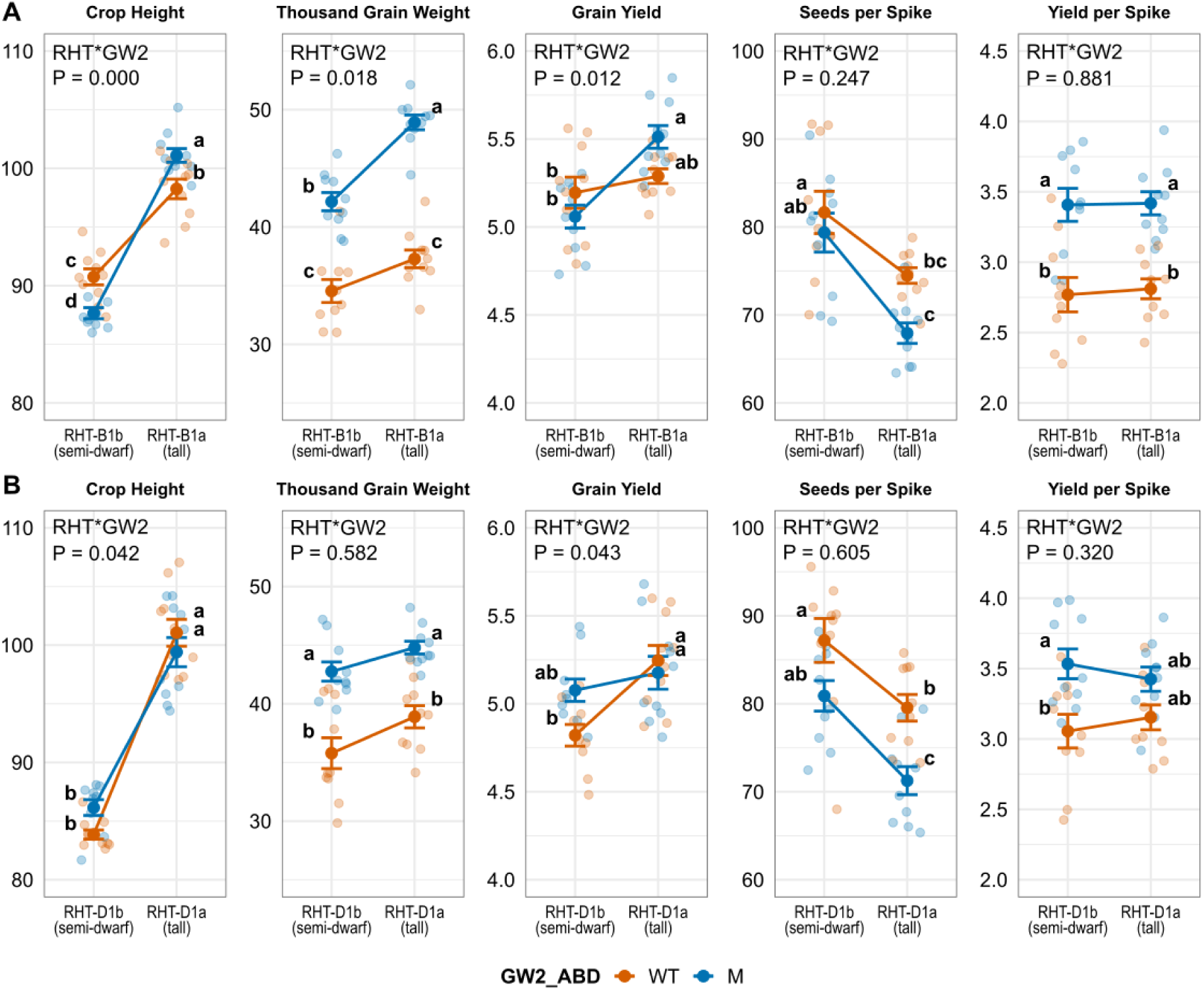
Interaction plots of selected traits for (A) BC_3_ *GW2*RHT-B1* NILs from and (B) BC_3_ *GW2*RHT-D1* NILs from Experiment 6. Wildtype NILs are shown in orange and the *GW2* triple mutant NILs in blue. Mean values (solid circles) are presented with standard error bars. Letters adjacent to each mean indicate groupings from post-hoc Tukey comparisons (α = 0.05) based on an ANOVA of combined genotype factor (*RHT Allele_GW2*). Interaction *p*values from a two-way ANOVA (*RHT Allele × GW2*) are displayed above each facet.

Significant interactions were also detected for grain yield, although these differed between *RHT-B1* and *RHT-D1* (Figures 4_A and 4_B). The *GW2* mutant yielded significantly less in the presence of *RHT-B1b*, whereas productivity was maintained with *RHT-D1b*. In both cases, the introduction of a semi-dwarfing allele reduced overall yield relative to WT (Supplementary Tables 8_A and 8_B).

No significant interactions were detected between *GW2* and *RHT1* for the two main spike yield components—seed number per spike (SSpk) and yield per spike (YldSpk). The expected trends were observed: semi-dwarfing *RHT1* alleles increased seed number per spike, while *GW2* mutants reduced it. Yield per spike was unaffected by *RHT1* but was elevated in the *GW2* mutant background.

Together, these results suggest that *GW2*-mediated increases in grain size are partially suppressed in the presence of semi-dwarfing *RHT1* alleles, particularly *RHT-B1b*. However, *RHT-D1b* did not negatively impact productivity in the *GW2* mutant background. This finding is of practical importance, as most UK cultivars utilise *RHT-D1b* to reduce plant stature.

### Secondary spike analysis confirms that spike yield is higher in all spikes (Experiment 6)

Phenotyping of main and secondary spikes, as described in Experiment 5, was repeated for the *RHT-B1b* (Figure 3_B; Supplementary Table 8_A) and *RHT-D1b* (Figure 3_C; Supplementary Table 8_B) NILs. Consistent with previous results, no significant differences were detected between main and secondary spikes when comparing *GW2* alleles. Thus, all spikes from *GW2* triple mutants contributed similarly to overall grain yield. Consequently, any reductions in total yield observed in these lines are likely attributable to decreased spike number per unit area rather than differences in per-spike productivity.

## Discussion

This study provides a comprehensive field-based evaluation of *GW2* in hexaploid wheat, combining multilocation, multiyear, and factorial experiments to elucidate its contributions to grain morphology, yield components and agronomic performance. Consistent with previous reports (Simmonds et al., 2016; Wang et al., 2018; Zhang et al., 2018), mutations in *GW2* homoeologues increased grain size and thousand grain weight (TGW) in an additive manner, confirming a conserved negative regulatory role for *GW2* in grain development in cereals (Kis et al., 2024; Li et al., 2010; Song et al., 2007). The single copy *A* genome mutant (*aaBBDD*) produced the most stable TGW gains among single mutants, while the combined triple mutant (*aabbdd*) achieved an average 20% increase in TGW across twelve field experiments. The stability of this effect across six breeder trials across a range of UK locations and sites underpins the robustness of *GW2* enhancement of grain weight. However, despite substantial increases in TGW, overall grain yield at the plot level was not significantly altered in the *GW2* triple mutant and was occasionally significantly decreased. This reflects the well-documented compensatory trade-off between grain size and grain number (Calderini et al., 1995; Fischer et al., 2014; Xie & Sparkes, 2021), where reduced grain number per spike and decreased spike numbers per unit area collectively offset the gains in individual grain weight. Comparable compensatory responses were also reported in a parallel set of field studies performed in Chile, where the *GW2* mutants displayed increased grain weight but reduced grain number across environments and sowing densities (Vicentin & Calderini, 2025).

Physiologically, this trade-off likely arises from overlapping developmental windows for grain number determination and ovary growth. *GW2* loss of function accelerates and enhances early carpel growth, increasing ovary weight during booting and anthesis (Simmonds et al., 2016; Vicentin et al., 2024). In doing so, it appears to limit the total number of developing grains, likely through altered assimilate partitioning or reduced floret fertility. Our data shows that, despite reductions in seed number per spike, yield per spike was consistently increased in the *GW2* triple mutant. This indicates that productivity at the spike level is enhanced; however, at the plant or canopy scale, overall grain yield remains constrained, presumably due to a reduced number of spikes per unit area. These findings are consistent with the “sink-limitation” hypothesis for wheat yield determination (Slafer et al., 2021; Slafer et al., 2023), which proposes that yield stability is maintained through developmental compensation among yield components. Supporting this interpretation, analysis of fertile spike number in our trials revealed the triple mutant produced approximately nine percent fewer spikes across Experiments 1–3. A lack of any genotype × environment interaction for final grain yield suggests that compensatory effects between grain number and weight in these lines occurs predictably, independent of growing conditions. Therefore, *GW2* represents a reliable modifier of grain size but is unlikely to drive yield improvements without additional genetic interventions that restore tiller number. Given the observed discrepancy between spike and canopy-level yield, we considered whether focusing solely on main tillers for spike yield component analysis may have biased results. However, evaluation of both main and secondary spikes in Experiments 5 and 6 showed that they responded similarly; higher TGW, fewer grains per spike, and increased spike yield. Although the magnitude of these effects was slightly smaller in secondary spikes, yield per spike remained consistently higher, indicating that the reduction in total grain yield observed in *GW2* mutants primarily reflects a lower number of tillers/spikes rather than reduced spike-level productivity. This interpretation is consistent with previous studies (Jian et al., 2024; Kis et al., 2024; Vicentin & Calderini, 2025) where *GW2* was also reported to play a role in regulating tiller number.

The stability of *GW2*-related phenotypes across extreme sowing densities (Experiment 5) suggests that the underlying developmental processes are largely independent of plant competition or tillering dynamics. Despite large differences in tiller number driven by seed rate, the relative increase in TGW and reduction in grain number were consistent, supporting the idea that *GW2* acts directly on grain formation rather than through alteration of assimilate competition among tillers or spikes. Although tiller number late in development was consistently lower in the triple mutant, the number of tillers per plant did not differ significantly, and the rate of tiller abortion was also comparable. This indicates that the lower spike number in *GW2* mutants likely results from reduced tiller initiation or poorer establishment, rather than increased tiller abortion or reduced tillering capacity per se. The observed reduction in plant establishment in the *GW2* triple mutant, particularly under high sowing densities supports this. Further study is required to determine the mechanistic effects of *GW2* on establishment and tillering dynamics. A clearer understanding of these subtle physiological effects and their contribution to reduced spike density will be essential for optimising yield performance in breeding programmes.

Given that the majority of modern wheat cultivars carry semi-dwarfing *RHT-B1b* and *RHTD1b* alleles, understanding their interaction with *GW2* is useful for breeding. Our data demonstrates that while *RHT* alleles predictably reduced plant height and TGW, *GW2* maintained its grain-size phenotype largely independent of *RHT* status. However, some allelic specificity was evident: *RHT-B1b* dampened the positive TGW effects of *GW2* and was associated with yield penalties in the *GW2* mutant background, whereas *RHT-D1b* did not significantly affect productivity. Importantly, discovering that the major positive effects of *GW2*, grain size and spike yield are compatible with *RHT-D1b*, suggests that introducing *GW2* mutations into modern semi-dwarf backgrounds should not compromise yield potential. This study highlights the inherent complexity of yield improvement via a single gene. While *GW2* loss of function consistently enhances grain weight and quality (including higher protein and gluten content), compensatory reductions in grain number and spike number prevent net yield gains. It should be recognised that the lines used here are backcrossed introgression lines derived from tetraploid *Triticum turgidum* cv Kronos and hexaploid *T. aestivum* cv Cadenza mutant populations into the Paragon background, and potential linkage drag may have influenced observed phenotypes. *GW2_A1* resides within a large haplotype block on chromosome 6A with limited recombination (Brinton et al., 2020), which could contribute to linked effects unrelated to *GW2* itself. Furthermore, these results were obtained within a single genetic background (cv. Paragon), and additional studies using alternative genotypes or breeding populations would confirm whether these trade-offs are conserved across different germplasm. Utilising targeted gene editing approaches could circumvent these issues by generating precise allelic variants directly within elite genetic backgrounds, thereby avoiding linkage drag and confounding background effects. Nevertheless, these mutants represent valuable genetic resources for breeding programs aiming to manipulate grain morphology.

Equally important is the validation of genetic effects under realistic field conditions. Robust, multi-year and multi-location trials are essential to accurately assess loci influencing yield and yield components. Controlled or single season experiments often fail to capture the full extent of genotype × environment interactions that determine agronomic performance. Multienvironment trials allows for the separation of genetic and environmental effects (Khaipho-Burch et al., 2023) and provides the most reliable measure of trait stability. This is especially critical for yield-associated genes, where compensation among components (grain number, grain weight, and spike number) can often either mask or amplify genetic effects depending on differing conditions. Through the UK Biotechnology and Biological Sciences Research Council (BBSRC) Designing Future Wheat (DFW) Breeders Toolkit programme, we accessed multi-location breeder trials that complemented our own experiments. The consistency of *GW2* phenotypes across multiple experiments, including the six breeder trials, provides strong evidence for its robust effects on grain morphology, and emphasises the need for validation prior to deployment in breeding programs.

Our results indicate that the *GW2-A1D1* double mutant offers a promising balance between enhanced grain size and limited yield penalty. Across experiments, this genotype produced consistent TGW gains (~14%) without significant reductions in yield, suggesting that partial suppression of *GW2* activity may be sufficient to achieve beneficial increases in grain weight while maintaining sink number. From a breeding perspective, targeting specific homoeologue combinations, particularly *A1D1,* could therefore enable introgression of grain-size improvements without compromising productivity. Furthermore, strategically combining *GW2* alleles with loci that increase spikelet fertility or floret survival e.g. *TaGNI1 (Sakuma et al., 2019)*, *TaCKX2.4 (Li et al., 2018)*, or tillering capacity e.g. *TaD27-B (Zhao et al., 2020),* may offset reductions in grain number. Future studies should focus on the physiological and molecular basis of this trade-off, including how different *GW2* dosage combinations influence assimilate partitioning and floret fertility. Integrating this knowledge with predictive breeding and genome editing will be key to unlocking the full potential of *GW2* in yield improvement.

## Conclusions

In summary, *TaGW2* acts additively across homoeologues to increase grain size and weight in wheat, and its effects are stable across environments, planting densities, and in the presence of *RHT-D1b*. However, complete loss of *GW2* function results in a consistent trade-off between grain size and number, leading to neutral or slightly negative effects on yield. The *TaGW2-A1D1* double mutant, by contrast, provides a valuable intermediate phenotype, combining increased TGW with maintained yield and may therefore represent the optimal target for breeding programs seeking to enhance grain size without incurring yield penalties. The absence of detrimental interactions between *GW2* and *RHT-D1b* further supports the feasibility of incorporating these alleles into semi-dwarf elite backgrounds. Realising yield benefits from *GW2* modification will likely require complementary alleles or agronomic interventions that restore spike number or enhance source capacity, thereby capitalizing on the larger sink potential conferred by *GW2* mutations.

## Acknowledgements

This work was supported the European Research Council (ERC-2019-COG-866328) and the UK Biotechnology and Biological Sciences Research Council (BBSRC) through the Designing Future Wheat (BB/P016855/1), Genes in the Environment (BB/P013511/1), Delivering Sustainable Wheat (BB/X011003/1), and Building Robustness in Crops (BB/X01102X/1) Institute Strategic Programmes.

## Author Contributions

**JS**: Conceptualization (equal); formal analysis (lead); investigation (lead); writing-original draft preparation (lead). **PC**: Investigation (supporting); writing-review and editing (supporting). **SE**: Investigation (supporting); writing-review and editing (supporting). **AML**: Investigation (supporting); writing-review and editing (supporting). **MK**: Investigation (supporting); writing-review and editing (supporting). **NB**: Investigation (supporting); writing-review and editing (supporting). **PT**: Investigation (supporting); writing-review and editing (supporting). **PJ**: Investigation (supporting); writing-review and editing (supporting). **DW**: Investigation (supporting); writing-review and editing (supporting). **CH**: Investigation (supporting); writing-review and editing (supporting). **DS**: Investigation (supporting); writing-review and editing (supporting). **CU**: Conceptualization (equal); funding acquisition (lead); writing-review and editing (lead).

## Supplementary Table Legends

**Supplementary Table 1**. Physical position and effect of the *GW2* mutations utilised in this study, sourced from the Kronos and Cadenza TILLING populations.

**Supplementary Table 2.** KASP markers and primer sequences used to screen germplasm utilised in this study.

**Supplementary Table 3.** A list of the field trials experiments performed and relevant information including year and location.

**Supplementary Table 4.** A list, description and trait ontology of the phenotypic assessments performed at each trial site.

**Supplementary Table 5.** Summary table of Paragon GW2 BC2/4 NILs evaluated in Experiments 1-3 (2019-2021). ANOVA and TUKEY analyses were performed by genotype + rep within individual years, and genotype + year (genotype*year) across years. The mean values per genotype, absolute differences Δ (delta), percentage difference from the WT (AABBDD) and Tukey’s HSD groupings (α = 0.05) are displayed for each trait. From the overall analysis the significance (P=) by genotype, year and interaction are also shown.

**Supplementary Table 6.** Summary table of GW2 BC4 Triple WT and triple mutant NILs in multi-site trials (Experiment 4; 2021). ANOVA was performed by genotype + rep for individual sites and genotype + location (genotype * location) across sites. Mean values per genotype, absolute differences Δ (delta), percentage differences and significance are shown for each location, with the genotype * location interaction also shown for the overall analysis.

**Supplementary Table 7.** Summary table of *GW2* BC_4_ NILs sown at three different seed rate (125, 275, 500 plants* m^−2^) in 2023 (Experiment 5). Analyses included (i) *Trait ~ Genotype + SeedRate + Rep + Stream + Genotype × SeedRate* to test main and interaction effects, and (ii) *Trait ~ Genotype_SR + Rep + Stream + Genotype_SR × Stream* to compare combined genotype–seed rate groups. The table reports mean values and Tukey’s HSD groupings (α = 0.05) for each genotype/seed rate combination, absolute differences Δ (delta) and percentage differences for genotype (WT / M), and p-values for genotype, seed rate and interaction.

**Supplementary Table 8.** Summary table of *GW2* x *RHT* BC_3_ NILs sown in 2023 (Experiment 6). Analysis of variance showing the effects of *Rhtb1* and *GW2* alleles and their interaction. Tukey’s HSD test was used to separate means, and letters indicate statistically significance (α = 0.05).

Results for the combined model are also provided, showing the overall effect of multiple allele combinations on trait performance. Mean values and Tukey’s HSD groupings (α = 0.05) for each *GW2/RHT* combination, absolute differences Δ (delta) and percentage differences are presented. *RHTB1 results are shown in A; RHTD1 results are shown in B*.

